# Expanding the epitranscriptomic RNA sequencing and modification mapping mass spectrometry toolbox with field asymmetric waveform ion mobility and electrochemical elution liquid chromatography

**DOI:** 10.1101/2022.10.28.514273

**Authors:** Richard Lauman, Hee Jong Kim, Lindsay K. Pino, Alessandro Scacchetti, Roberto Bonasio, Benjamin A. Garcia

## Abstract

Post-transcriptional modifications of RNA strongly influence RNA structure and function. Recent advances in RNA sequencing and mass spectrometry (MS) methods have identified over 140 of these modifications on a wide variety of RNA species. Most next-generation sequencing approaches can only map one RNA modification at a time, and while MS can assign multiple modifications simultaneously in an unbiased manner, MS cannot accurately catalog and assign RNA modifications in complex biological samples due to limitations in fragment length and coverage depth. Thus, a facile method to identify novel RNA modifications while simultaneously locating them in the context of their RNA sequences is still lacking. We combined two orthogonal modes of RNA ion separation before mass-spectrometry identification: high-field asymmetric ion mobility separation (FAIMS) and electrochemically modulated liquid chromatography (EMLC). FAIMS RNA-MS increases both coverage and throughput, while the EMLC LC-MS orthogonally separates RNA of different length and charge. The combination of the two methods offers a broadly applicable platform to improve length and depth of MS-based RNA sequencing while providing contextual access to the analysis of RNA modifications.

## Introduction

Since their initial discovery in the 1950s(1), RNA modifications have recently returned into interest due to advancement of specialized next-generation sequencing (NGS) and mass spectrometry (MS) methods, which have led to the identification of over 140 covalent modifications of RNA to date and to genome-wide mapping of a small subset of them (2–5). Recent examples of the biological impact of RNA modifications can be found in the discovery of tissue-specific patterns of N^6^-methyladenosine (m^6^A) on mRNA and the importance of 7-methylguanosine (m^7^G) in microRNA biogenesis (7,8). While neither NGS nor MS alone can identify the nature and position of all existing RNA modifications, both techniques used in tandem can provide orthogonal perspectives.

Currently, most NGS approaches to detect RNA modifications rely on antibodies or enzymatic/chemical treatments and carry some limitations: 1) the availability and specificity of certain antibodies/treatments can create obstacles to accurate mapping of the modification, 2) only a single modification can be measured at a time (9). Nanopore sequencing has demonstrated base resolution of m^6^A but has yet to be adapted to other types of RNA modifications (7,10). On the other hand, RNA MS provides an unbiased approach to the identification, quantification, and mapping of a large variety of RNA modifications; however, the method is currently limited by the availability of amenable liquid chromatography strategies to reduce the complexity of highly modified and complex ribooligonucleotide species (3,11). Given that RNA contains a highly negatively charged phosphate backbone and polar bases, the traditional forms of chromatography that are typically paired with MS, such as reverse-phase resins and mobile phases developed primarily for peptide separation, do not effectively retain RNA(12,13). Several methods are being developed to circumvent this obstacle, including using suppressive ion pairing reagents, but all available strategies have severe drawbacks, especially in co-opting on instruments also utilized for peptide detection (13–15).

Typically, reverse phase high pressure nano liquid chromatography (HPLC/nanoLC) is utilized to reduce complexity of peptide mixtures injected in the MS in each time window. However, even HPLC is limited by complexity when selecting low-abundance species in an elution window. To overcome such limitations, various strategies have been devised. One example is ion mobility mass spectrometry (IM-MS), where ions are selected in the gas phase depending on how their specific charge density interacts with oscillating voltage fields: their separation is determined by the cross-sectional area of the molecule (16–18). A specific form of IM-MS utilizes high-field asymmetric waveforms (FAIMS), also known as differential mobility spectrometry (DMS). In two studies, this approach improved selection of proteins for top-down fragmentation and increased depth of peptide proteome, but it has never been applied to ribooligonucleotide MS (16,19,20). A major advantage of ion mobility/DMS, such as FAIMS, is the orthogonality to liquid chromatography separation of complex mixtures or isobaric species, such as modified RNA bases (20,21). To improve separation and selection of both modified nucleosides and ribooligonucleotides, ion mobility MS has been successfully applied to multiple biological systems and molecules, however, the use of ion mobility to distinguish diverse ribonucleotides and their modifications has been limited, but no one has applied FAIMS (DMS) to sequencing RNA by MS (21–23). More importantly, the use of traditional ion mobility approaches requires a considerable amount of expertise and specific MS instrumentation to design new platforms, from MS acquisition to data processing. Thus, intuitive and robust methods must be designed to improve and expand the limited toolbox for RNA by LC-MS/MS.

In this study, we developed a new FAIMS method for RNA MS and a new voltage-based chromatography method for ion pairing free RNA LC-MS/MS. Our work synergistically combines two new approaches rarely used for RNA MS analysis into a single platform: 1) an ultrafast 2-minute FAIMS MS RNA sequencing method and 2) a new form of RNA chromatography for increased separation independent of reverse phase resins. Utilizing both direct injection and electrochemically modulated liquid chromatography (EMLC) coupled with FAIMS, we can sequence modified RNA molecules to high accuracy. Our method enhances quality and speed of data acquisition for RNA MS and thus create opportunities for new experiments for characterization of RNA.

## Methods

### Human rRNA Extraction and Digestion

HeLa cells were grown in Thermo Fisher DMEM (#12491023) supplemented with 10% fetal bovine serum albumin (#16000044) with 1x Thermo Fisher Pen/Strep (#15140122) antibiotics in quadruplicate 10cm plates. Plates were scraped and lysed using the QIAGEN RNeasy (#74106) protocol to extract total RNA. Total RNA (1ug) was then fragmented using NEB Mg (II) (#B9021S) RNA fragmentation for 10 minutes at 95°C to randomly fragment RNA species. The RNA species or mixture was then cleaned by Glygen strong anion exchange TopTips™ to desalt.

### Direct Injection RNA FAIMS

The *E coli* and *S. cerevisiae* tRNA standard was purchased from Thermo Fisher and digested with both RNase T1 and RNase A (#T3018) from NEB biosciences for 45 minutes at room temperature in triplicate. The *Let7a* microRNA standard was purchased from IDT using the normal desalting measures. The RNA species or mixture was then cleaned by Glygen strong anion exchange TopTips™ to desalt. The RNA was dried in a Thermo Scientific Savant™ SpeedVac™ and reconstituted into 30% LC-MS grade acetonitrile (#047138.K2), 30% methanol (#615130025) and 1mM ammonium formate (#014517.30) at pH 5. The Advion Nanomate™ was set to default settings and a spray setting of 1.9kV and waiting for 5-10 seconds for spray stability before running with a run time of one minute each run. Each technical replicate is an injection of the digested tRNA into the Thermo Scientific Fusion™ MS and sprayed using the default flow settings for 2 minutes per injection at a concentration of 200 ng/μL (at 10 μM for *Let7a*). The ionization spray voltage was kept steady at 2.2 kV in negative mode and with the Thermo Scientific FAIMS Pro II attached we had no gas flow and “User Settings” of 70/70°C for the inner and outer electrode temperatures. The compensation voltages (CV) used in the direct injected experiments were 30, 40, 50, 60, 70V. The MS^1^ maximum injection time (MIT) was set to 100ms and an AGC target of “Standard” for both the FAIMS attached and No FAIMS (NF) and a HCD normalized fragmentation of 20% for all oligonucleotides of all experiments. Species were chosen for MS^2^ by being greater than a charge state of 3 and a minimum intensity of 1×10^5^. The MS^2^ spectra was acquired using a 200% AGC target and a 300ms MIT for both direct injection and EMLC.

### Electrochemically Modulated Liquid Chromatography

The PGC (loose resin from 35003-031030) EMLC used a Thermo Easy™ 1000 nano LC with mobile phase A consisting of 5mM ammonium formate at pH 5 and mobile phase B consisting of 30% acetonitrile, 30% methanol and 5mM ammonium formate at pH 5. The voltages applied to the in-house fritted and packed PGC trap column were generated from a Volteq HY3006D voltage power supply instrument with a programmed (available on GitHub https://github.com/heejongkim/voltageGardient) gradient applied in a stepwise increase throughout the run, increasing with the linear gradient of mobile phase B. A grounding wire was placed between the voltage trap column and the tip to ensure source voltage does not affect the column. The MS method was the same as with or without the FAIMS cycling through the same previously used 40 to 70 CV values throughout the 90-minute total gradient time. The linear hydrophobic gradient started at 0% from 0-10 minutes, then to 30% at 50 minutes, then 60% at 70 minutes followed by a wash.

The acquired data for both method types was processed and visualized using the OPEN MS platform 2.6.0 (TOPPAS and TOPPView) and the NASE RNA sequencing software (24). The Savitsky-Golay Noise Filter and Baseline filter modules were used to better optimize our data processing before the use of NASE. Our settings for NASE were 10ppm MS^1^ and 10ppm MS^2^ tolerances, salt adducts of Na(+), K(+), and Mg(2+) were used, and all possible RNA fragment ions (a,b,c,d and w,x,y,z). The use of the a-Base fragment ion was used to resolve the base and 2-O-methyl modification when the spectra contained sufficient ion populations. The minimum size of the RNA was at 5 nucleotides with a maximum of 25 and a missed cleavage at 3 for RNase T1 or RNase A. Gene sequences for both *S. cerevisiae* and *E. coli* were taken from the Genomic tRNA Database and sequences were shuffled to create the decoy databases (25). Given the limited number of spectra, an FDR of 10% was used with a hyperscore 40 score cut off for modifications in both the direct injection and chromatography acquisitions. Figures were rendered using ggplot package with the rawDiag package and base R functions (26). All data uploaded to PRIDE under the reference [1-20220918-50331].

## Results

### Development of an Ultra-Fast Ion Mobility RNA MS Method with FAIMS

Our work began with a simplified approach to RNA MS by removing the LC instrumentation and solely utilizing direct injection MS analysis to ascertain if FAIMS can improve RNA modification coverage. FAIMS ion selection is achieved by an asymmetric alternating voltage, named the compensation voltage (CV; also known as the dispersion voltage, DV), using high or low asymmetric field applied between an inner and outer electrode selects for analytes or populations of analytes (27). To increase the depth of coverage of RNA MS with FAIMS, we first coupled a Nanomate™ direct injection electrospray system to a Thermo Orbitrap Fusion™ instrument configured with FAIMS and tested the lower limits of FAIMS selection with a synthetic 3’ biotinylated *D. melanogaster* 22nt *Let7a* microRNA (UGAGGUAGUAGGUUGUAUAGUU) used previously in our lab as a standard (Fig. S1A) (24). With no FAIMS installed (NF; Fig. S1A, top) the mass spectrometric MS^1^ signal from the microRNA was lost within the background of singly charged contaminant species, but using FAIMS (Fig. S1A, bottom) the microRNA signal was enriched over the background with multiple charge states identified as calculated by MongoOligo electrospray series (5).

To further test the selective power of FAIMS, we focused on tRNAs, a highly modified group of RNAs known to be difficult to sequence by NGS (28,29). We infused a mixture of RNase T1 digested *E. coli* tRNA (Fig. 1A). Ramping the FAIMS compensation voltage (CV) from 30 to 70 CV (in increments of 10 CV) selected for a different distribution of digested RNA species (shown by intact mass) in comparison to no FAIMS (NF) (Fig. 1B). The distribution of species enriched by each CV ranged from 6 nts (left dotted line on axis) selected at 70 CV to 20 nts and above in the 40 CV (right dotted line on axis), calculated assuming a non-modified RNA species. Mirroring this result, analyte charge states decreased with longer selected RNA species, with charge states decreasing through the CV cycle (Fig. 1C). These observations indicate that FAIMS can separate RNA species by charge, and, in turn, by size, given that charge is correlated to RNA length due to the phosphate backbone (30).

**Figure 1.**
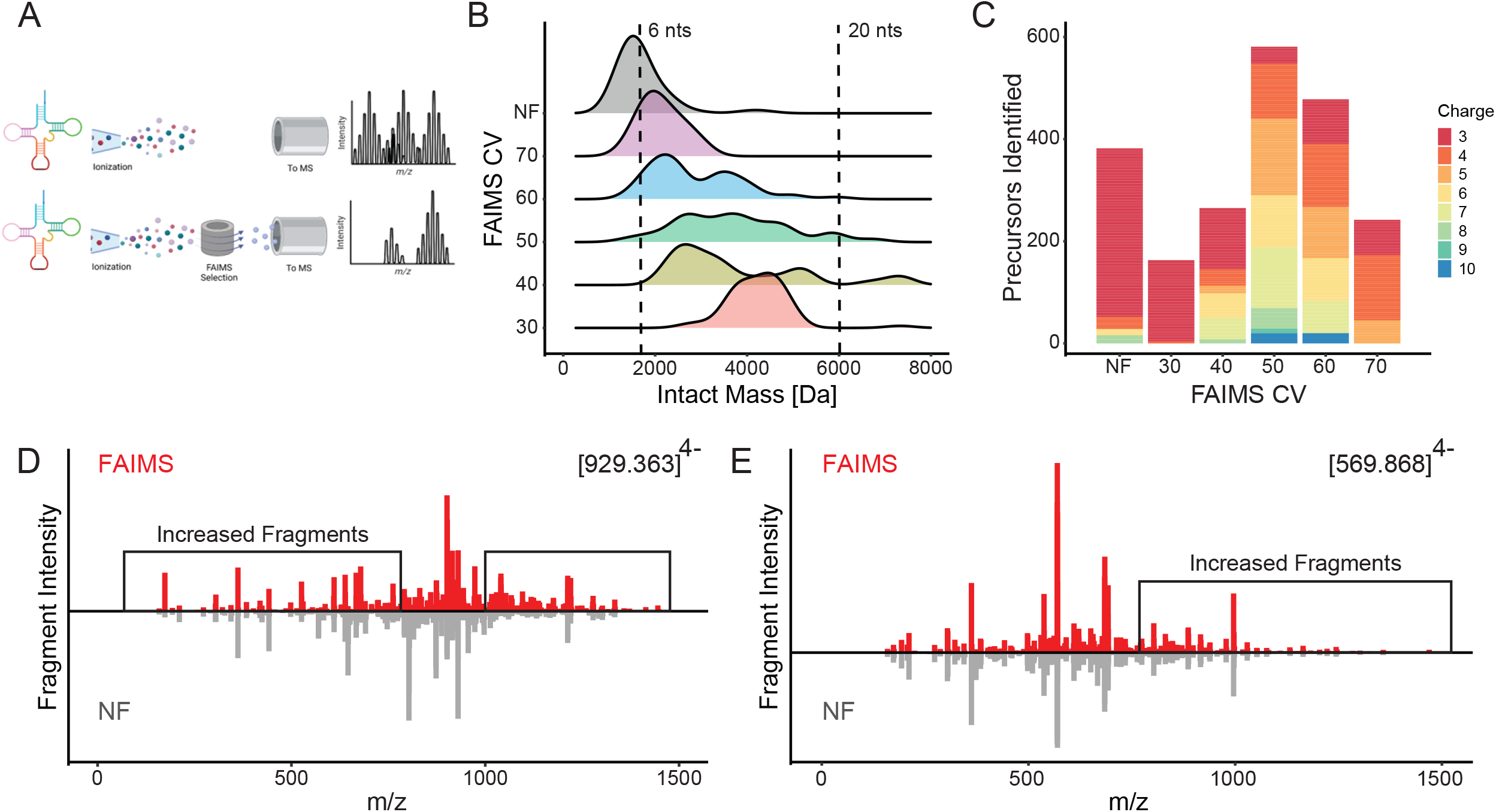
Direct Injection FAIMS RNA MS. A) Diagram of FAIMS selecting for individual tRNA species. B) Distribution of E. Coli tRNA digested fragment sizes shown in intact mass [Da] with FAIMS CV selection in comparison to FAIMS not attached (NA) comparing two single injections with dotted lines illustrating 6nts (left) and 20 nts (right) C) Charge state distribution of each CV in comparison to FAIMS not attached (NA) using the same two single injections of tRNA D) Sequenced *E. coli* tRNA oligomer UCCCCCCCCUCGp compared between FAIMS (top) and NA (bottom) E) Sequenced *E. coli* tRNA oligomer UCCACUCAGp compared between AIMS (top) and NA (bottom)

This increased separation of diverse RNA species, however, doesn’t necessarily correlate with increased mass spectra quality. To directly compare spectral quality from FAIMS to NF, the two highest confidence fragmentation sequencing spectra (MS^2^) in the NF dataset, assigned to RNA sequences UCCCCCCCCUCGp (Fig. 1D, Fig. S1B) and UCCACUCAGp (Fig.1E, Fig. S1C) were overlayed with the spectra obtained for the same RNA oligonucleotides using FAIMS. The spectra for UCCCCCCCCUCGp acquired with FAIMS (Fig. 1D, red) showed an increased intensity and captured a greater variety of RNA MS^2^ fragments and led to an overall higher confidence in their interpretation compared to the spectra obtained without FAIMS (NF) (Fig. 1D, gray). A similar improvement in spectra quality when using FAIMS was observed for UCCACUCAGp with both increased MS^2^ fragments and increased intensity (Fig. 1E). Thus, each FAIMS CV selects for a unique distribution of RNA species and the increased selectivity results in improved tandem mass spectral quality.

### Increased tRNA sequence and modification coverage with FAIMS

We next sought to determine whether FAIMS can increase the confidence of modification assignment and coverage of RNA species. We again analyzed an E. coli tRNAs digested mixture (See Methods) while attempting to simultaneously determine their sequence and by identifying and localizing chemical modifications with and without the use of FAIMS. Peptide sequencing by mass spectrometry has utilized hyperscores (hscores) to determine the confidence in each observed peptide by comparison to an *in silico* predicted peptide MS^2^ fragmentation spectra(31). Similarly, for RNA MS/MS, hscores can be utilized to gauge the quality of ribooligonucleotide spectra, as shown for our previously published program Nucleic Acid Search Engine (NASE) (24,32). Hscores from FAIMS and NF combined triplicate injections sequenced using NASE demonstrated a statistically significant (*p* = 1 × 10^−17^, Wilcoxon Rank Sum) increase in spectra confidence with FAIMS (Fig. 2A). The improved number and confidence of MS^2^ fragmentation spectra can be attributed to the selection of FAIMS, which reduces background ions and increases the population of informative RNA-derived species, decreasing ion fill time (injection time) (Fig. S2A) (33,34). This result demonstrates the potential of FAIMS beyond the separation of RNA ions in leading to increased duty cycle time to sequence and identify additional RNA species.

**Figure 2.**
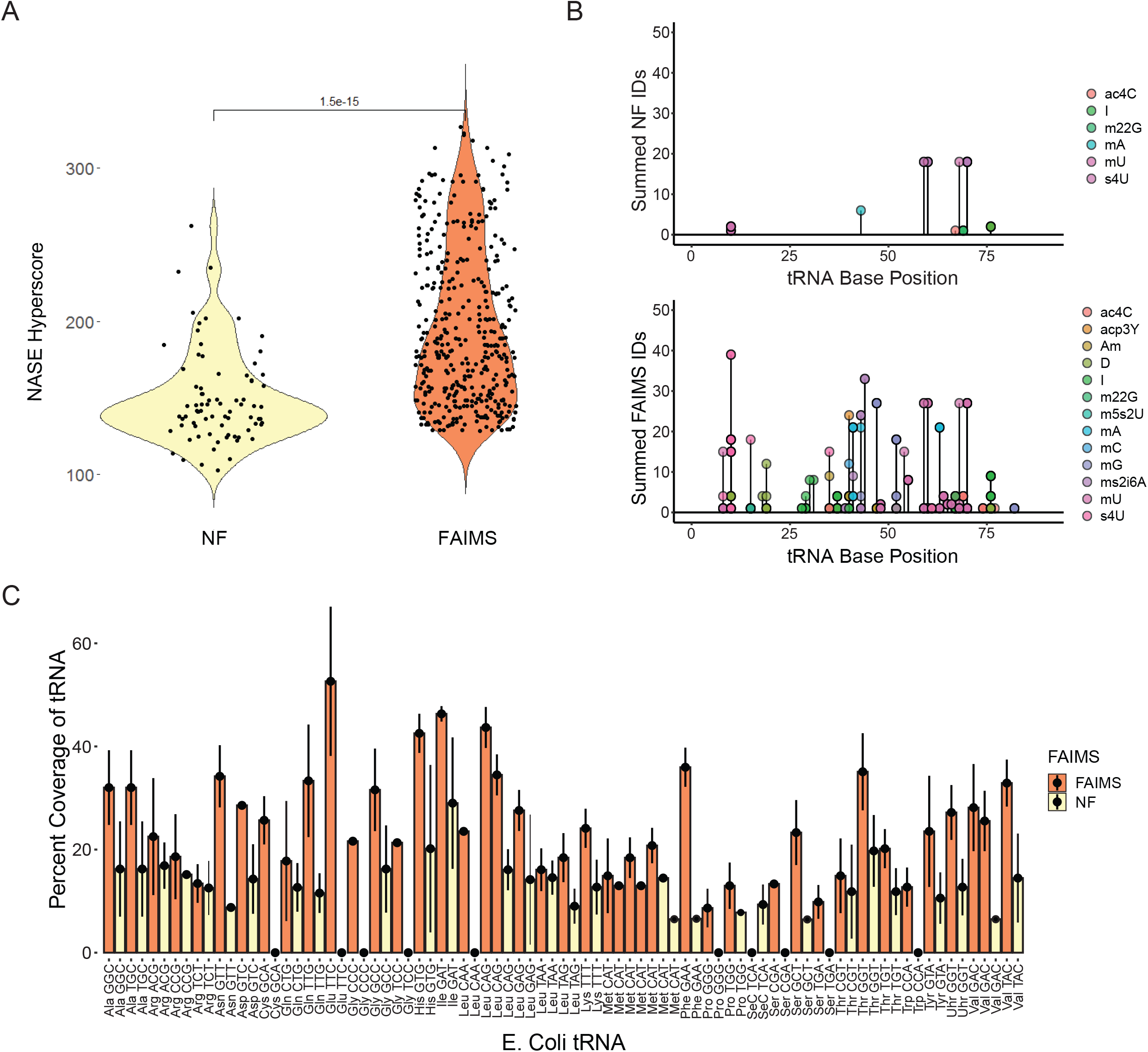
Differences in Coverage and Modification Identifications using FAIMS A) Improvement of NASE hyperscore sequenced spectra of E. coli tRNA sequences in triplicate runs for both comparing with FAIMS and without (NF) using a Wilcoxon Rank sum for statistical significance. B) Modification coverage of *E. Coli* tRNA with the sequenced RNA species colored in both NF(top) and FAIMS(bottom) with species checked for accuracy from the Modomics database. C) Comparing FAIMS and no FAIMS (NF) over triplicate runs for summed total coverage separated by tRNA gene.

Having demonstrated that FAIMS provides higher-confidence spectra, we next proceeded to analyze the chemical modifications in *E. coli* tRNA. tRNAs are highly modified and thus served as an ideal study case for this purpose (35). We performed a database search of the most common *E. coli* tRNA modifications and found that our ability to detect them was dramatically increased after differential ion selection via FAIMS (Fig. 2B) (36,37). The FAIMS characterized RNA modifications align with previously published data. For example, we observed dihydrouridine (D) within the D loop of tRNA (30–35 base position), consistent with previous studies, validating our new MS sequencing technique(36). Overall, compared to the MS in the standard configuration, FAIMS MS achieved on average a two-fold coverage increase in sequence coverage than NF across all *E. coli* tRNA genes (of a total of 72 unique *E. coli* tRNAs identified), eight of which were only detected with FAIMS (Fig. 2C). Even with the high homology found between tRNA sequences(38), our direct injection FAIMS-based mode was able to sequence a minimum of two unique oligonucleotide spectra per tRNA species (Fig. S2B). Thus, we have utilized FAIMS to increase the number of identifications, with increased quality of spectra, resulting in increased coverage across all tRNA species. In addition, to increase tRNA coverage beyond RNase T1 fragments, we digested with RNase A. While RNase T1 total IDs were greater than RNase A total IDs (Fig. S2C–D), the complementary digestion allowed for increased diversity of RNA species and increased coverage of the 5’ region of the tRNA (Fig. S2E).

Overall, the use of FAIMS for direct inject ultra-fast identification of RNA modifications by MS and increased coverage of tRNA is vastly superior to acquiring samples without FAIMS and is compatible with two common types of RNA digestions. This work stands to further expand the toolbox for RNA analysis by MS. Several advantages are obtained with FAIMS by limiting the amount of background ions, while also identifying the most abundant modifications on highly modified tRNA.

### Electrochemical Modulation Liquid Chromatography for separating RNA by nano LC-MS/MS

While direct spray infusion is higher throughput, it is also inherently limited on coverage depth (even with the addition of FAIMS). Current liquid chromatography separations of RNA are challenging, as they often require LC conditions (ion pairing reagents) which are not compatible with the typical MS instrument laboratory. To overcome the difficulties, we developed a new RNA LC-MS/MS method based porous graphitic carbon (PGC), which is commonly used to retain polar and hydrophobic compounds due to the aromatic structure and varied sizes (39). We took advantage of the conductive nature of PGC to retain the RNA through base stacking and polarity, similar to the other electrochemically modulated liquid chromatography (EMLC) designs reported previously (Fig. 3A) (39–41). To test the column retention of RNA to conductive PGC, we extracted total RNA from HeLa cells and fragmented it by heating in presence of Mg^2+^, to increase the size diversity of RNA oligonucleotides (42,43). The EMLC gradient was composed of both reversed phase liquid chromatography and voltage changes. First, voltages were stepped in discrete values from 0 to 25V with a short linear hydrophobic gradient utilized for each voltage step. Increasing voltage provided a different elution profile (as shown through MS^2^ acquisitions) throughout the run, which illustrated the differing mobility of RNA with increasing voltages (Fig. 3B). The length of the RNA fragments played a major role in retention times and their response to increasing voltages with the same linear LC gradient for each voltage. Different RNA length profiles eluted at each step until 25V were reached, indicating that 30V was sufficient for the elution of longer, more informative RNA species (Fig. 3C). Until 25V, each increasing voltage released longer RNA or a more diverse set of RNA species; however, multiple individual voltage gradient LC-MS/MS acquisition to separate total RNA are time-consuming and the total run time could be reduced by combining the voltages together.

**Figure 3.**
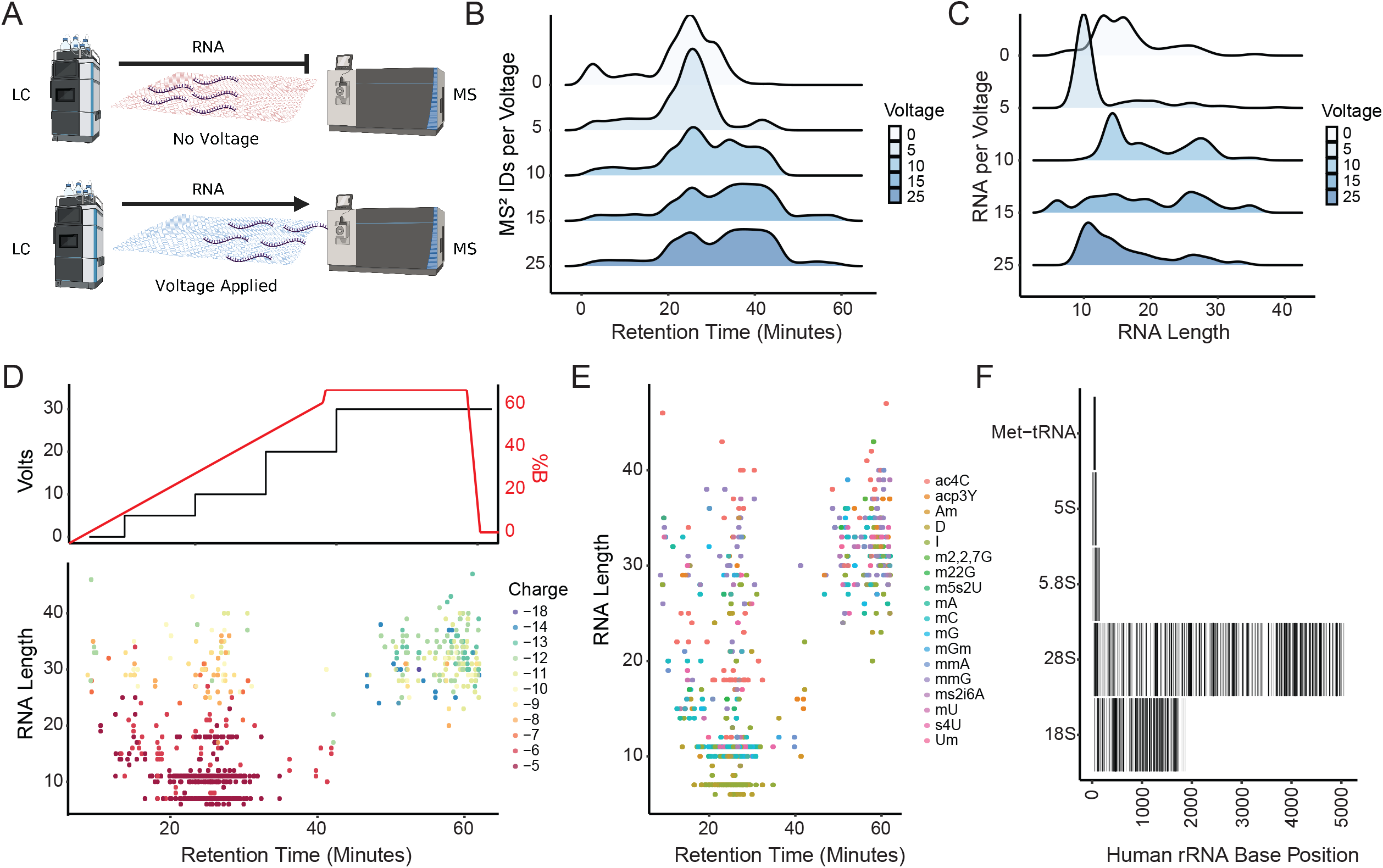
Separation of Digested rRNA and Identification of Modifications using EMLC. A) Schematic for the EMLC gradient system. B) Populations of spectra (MS^2^) identifications taken with increasing static column voltages during a reverse phase gradient. C) Populations of RNA length elutions with increasing static column voltages during a reverse phase gradient. D) Stepwise votlage elution with reverse phase with voltages plotted (top) and RNA length and charge state plotted to elution (bottom). E) Modifications overlaid onto RNA length elution over the EMLC gradient. F) Total coverage of the human 80S ribosome RNA distributed over the known base positions using the sum of 4 replicates.

To elute all the RNA oligonucleotides within a single gradient, as occurs in LC-MS for peptide analysis, we combined a stepwise voltage gradient from 0 V to 30 V with a linear hydrophobic gradient (Fig. 3D top). In the first 30 minutes of this combined chromatographic method, the identified RNA species were predominately short (less than 10 nts) with low charge states (greater than -6), however, the longer fragments eluted from the column considerably later (Fig. 3D, bottom). Even RNA of similar length and lower charge elute earlier than the highly charged species of the same length, leading to the hypothesis that charge drives elution. As modifications can mask or influence charge and structure of RNA species, we examined where modified RNA species are preferentially retained. Modified RNA showed no selectivity in elution, with the same known modifications eluting throughout the gradient with parity (Fig. 3E) (43). In the total identified sequences, we uniquely mapped 73% of the 18S and 65% 28S ribosomal subunits (Fig. 3F). Thus, our modified electrochemical liquid chromatography approach improves the ability to handle ion complexity and is amenable for any LC-MS/MS platform.

### Combining FAIMS and EMLC RNA MS Platform

We reasoned that combining separation steps based on ion separation in both liquid phase (EMLC) and gas phase (FAIMS) would synergistically increase our ability to deconvolute complex RNA mixtures. Next, we combined *E. coli* and *S. cerevisiae* tRNAs, digested with RNase A and RNase T1 for 30 minutes and analyzed them using FAIMS with or without EMLC. First, the addition of EMLC increased the RNA MS/MS spectra identifications over 500-fold against direct injection FAIMS and resulted a drastic improvement over the EMLC alone *E. coli* and *S. cerevisiae* tRNA (Fig. 4A). As expected overall the use of a gradient to further deconvolute the mixture increased the number of spectra acquired and improved total identifications and sequence coverage with the combined use of FAIMS and EMLC. The total number of spectra per tRNA is improved to the same level as the total spectra acquired per run, with an equally improved coverage across tRNA in comparison to FAIMS and EMLC alone (Fig. 4B). The number of modifications for both the *E. coli* and *S. cerevisiae* tRNA increased both coverage and identifications (Fig. 4C). The summed modifications of both tRNA subsets increased with the FAIMS EMLC design, dramatically improving the coverage of the modified regions. Again, we identified the characteristic D-loop modifications with increased spectral counts per identified site localized sequence in comparison to FAIMS without LC. Overall, the synergistic use of both EMLC and FAIMS increases the sensitivity and coverage of modified RNA species.

**Figure 4.**
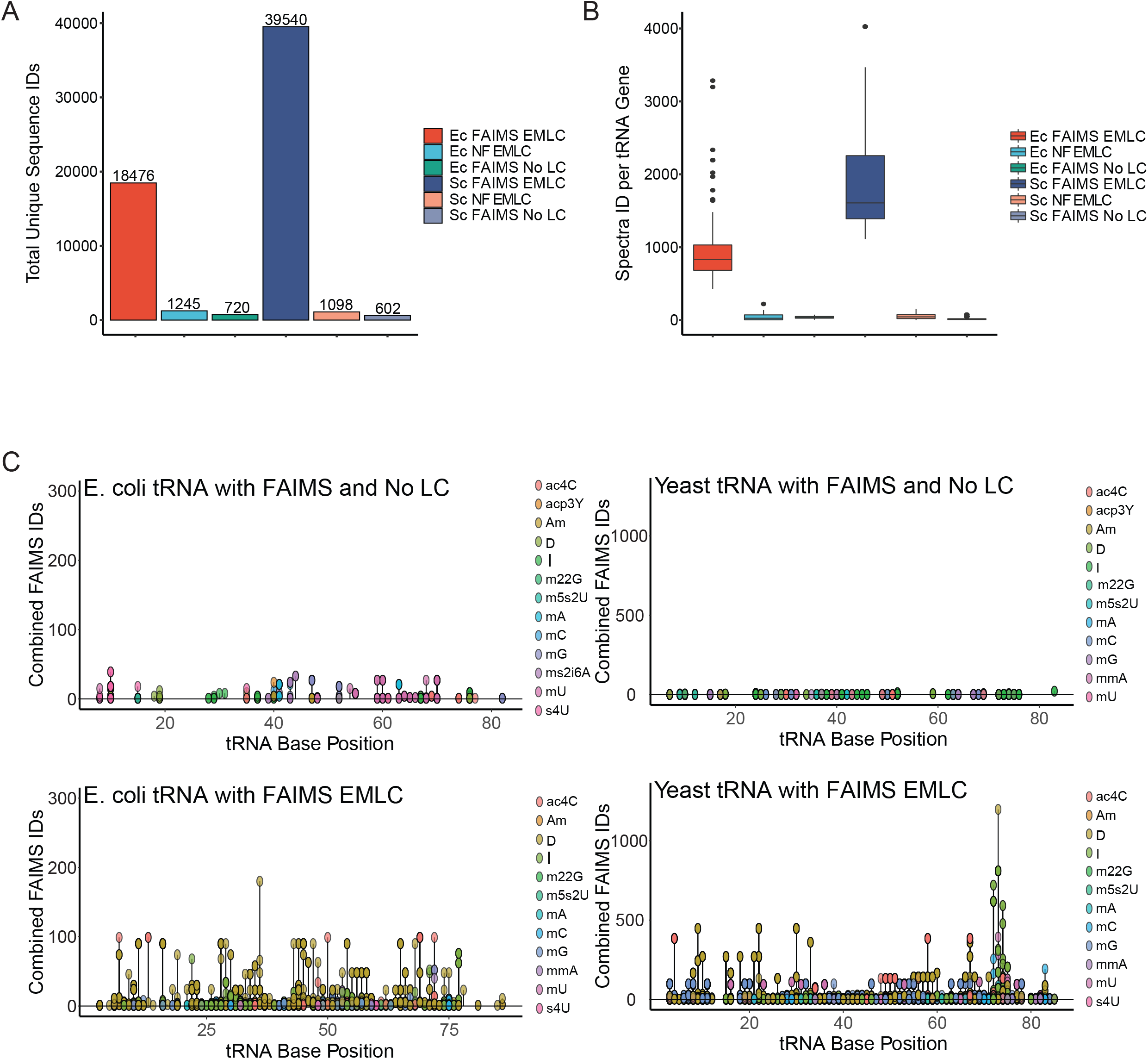
The combination of FAIMS and EMLC to sequence E. coli and Yeast tRNA modifications. A) The total unique sequence IDs (unique tRNA sequence spectra) for FAIMS EMLC, No FAIMS (NF) EMLC, and Direct Injection FAIMS (FAIMS No LC) from Fig.2B for both E. coli (Ec) and S. cerevisiae (Sc) samples. B) Distribution of sequenced spectra per tRNA gene for the same samples outlined in A. C) Linear plots of summed modification IDs across all tRNA (top left the same as Fig.2B) distributed by method and species with ID axis normalized to their respective species.

## Discussion

We have developed a new method to improve the speed of detection, sensitivity and coverage of RNA modifications through increased retention and orthogonal separation of RNA. Each CV value selects a different pool of RNA molecules, many of which would not be readily identifiable in a global mixture, thus providing a means for larger scale transcriptomic analyses by LC-MS/MS when FAIMS is used in combination with EMLC. Our work improves upon the impressive recent development of a large-scale RNA sequencing algorithm, the original design of FAIMS, as well as existing ion mobility approaches for RNA MS analysis (21,24,27).

In addition to applying FAIMS to RNA MS for the first time, we applied the principle of EMLC to further improve RNA fragment separation before MS(21,40,44). We developed a combined voltage and chemical gradient that effectively separates RNA fragments by size and charge (Fig. 3). This gradient design selects RNA through charge-based interactions with the stationary phase (PGC) and as the stationary phase charge retention changes, so does the retention of RNA. Our hypothesis is that the dominant phosphate charges of the RNA backbone interact with the induced dipole of the PGC and that, as the voltage is increased, the strength of these interactions is reduced resulting in a gradual release of the RNA fragments(39).

Overall, the combination of these two methods has created a new platform for sequencing RNA modifications using mass spectrometry. One of the major advantages of RNA MS is the discrete identification of multiple modifications simultaneously, however our improved method allowed to locate their position within an RNA molecule. This is crucial for highly modified species of RNA, such as tRNA, in which densely modified regions require site specific localization to deconvolute. The ability to site localize and concretely distinguish RNA modifications on multiple different overlapping RNA species provides a crucial orthogonal view to NGS sequencing, while also performing an unbiased and far deeper RNA modification coverage with FAIMS. Beyond this, the method reduces the number of “contaminant” species present in the spectra, therefore wasting less time on smaller, non-unique RNA or non-RNA species. Finally, the use of two method designs generated a more high-throughput MS approach for the quantitation of RNA modifications or longer LC-based gradients for increased depth of sequence and modification coverage. This new method is an improvement over the previous designs and is amenable to almost any RNA-MS platform.

These new techniques expand the toolbox for the rapidly expanding field of epitranscriptomics. It enables LC-MS/MS users to discover and quantify RNA modifications with an orthogonal approach to common NGS methods. In the future, we envision that the study of RNA modifications will take advantage of a blend of NGS and LC-MS/MS techniques, each with complementing strengths. For MS, we must further develop methods to increase the length of RNA sequenced and improve statistical confidence in site localization of post-transcriptional modifications. We believe our work provides an essential step forward toward the goal of comprehensive and quantitative mapping of the epitranscriptome.

## ASSOCIATED CONTENT

### Supporting Information

All raw datafiles and processed files are available from download from PRIDE under the reference number [1-20220918-50331]

The Supporting Information is available free of charge on the ACS Publications website.

Comparison of both FAIMS and NF oligonucleotides (.xlsx)

## AUTHOR INFORMATION

### Author Contributions

The manuscript was written through contributions of all authors. All authors have given approval to the final version of the manuscript.

## Acknowledgements

We thank the Weitzman laboratory for their focused and critical feedback on the figures and written manuscript. We would like to thank Scott Peterman and Mike Belford of Thermo Scientific for their feedback on the use of FAIMS. We humbly thank both Josue Baeza and Stephanie Lehman Miller for their consistent support and insightful feedback on the project. Funding from NIH grants CA196539, AI118891 and HD106051 to BAG and R01GM127408, R01GM138788 to RB are gratefully acknowledged.

## Table of Contents Art

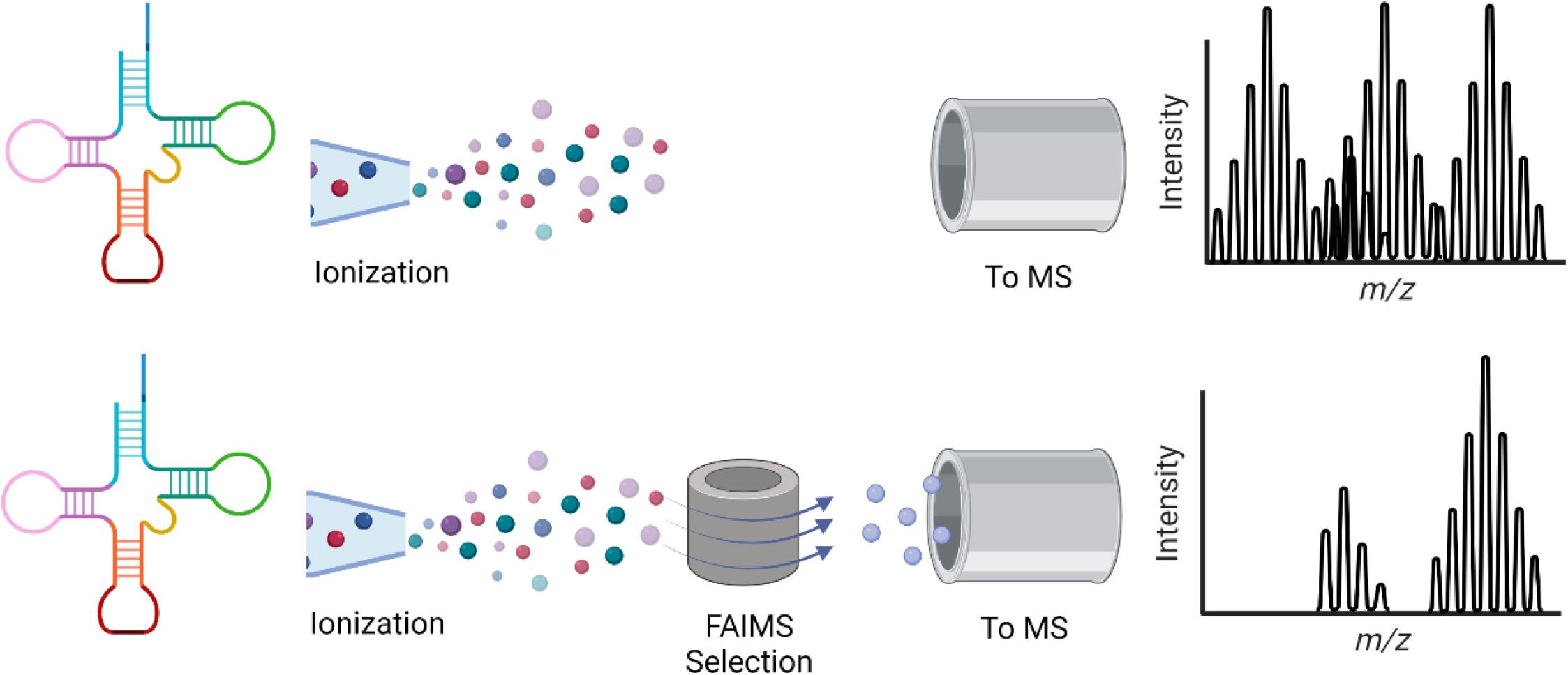

---

**Supplemental Figure 1.**
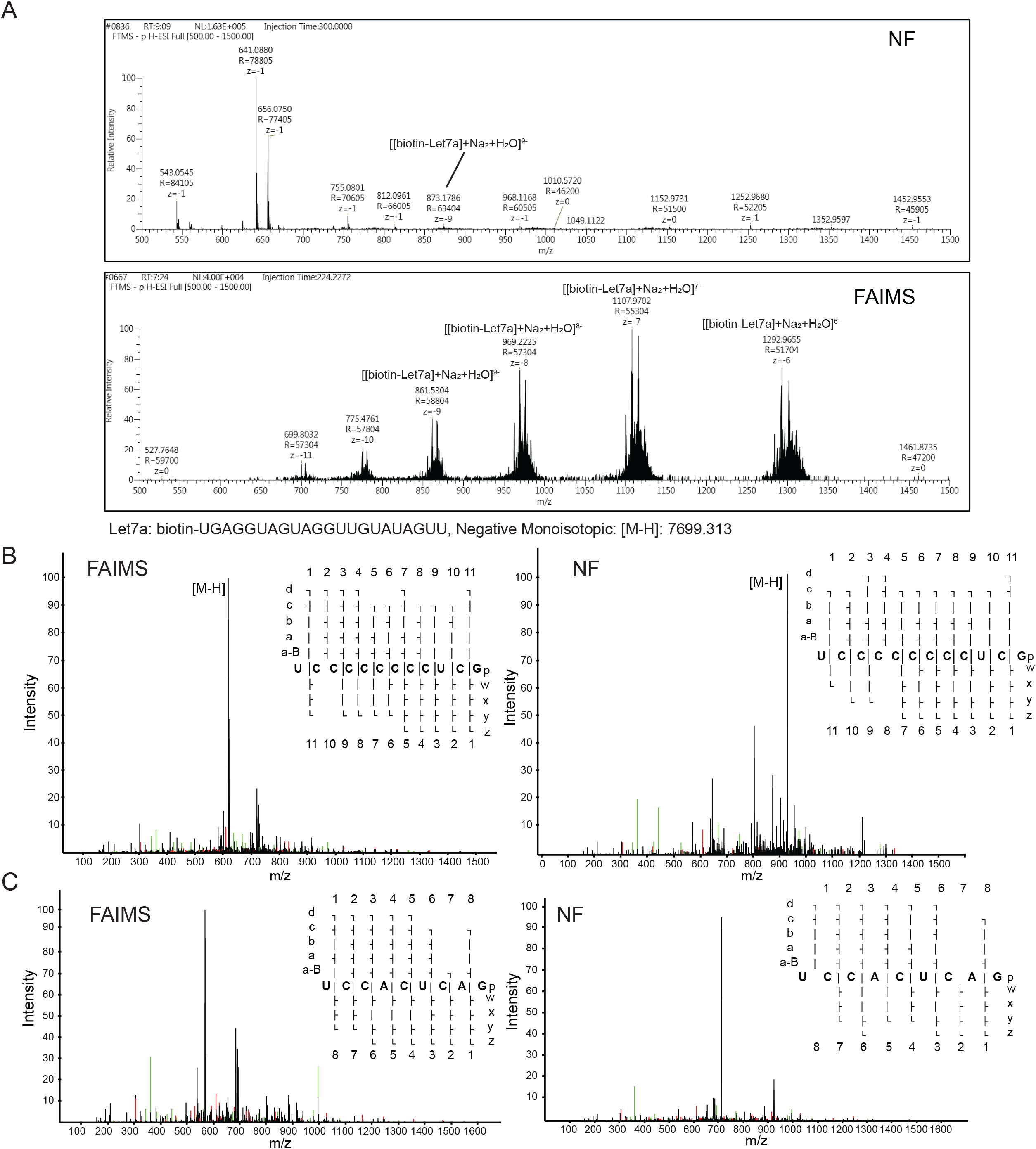
Direct Injection FAIMS Let7 microRNA MS. A) Spectra of 10nM Let7a microRNA (UGAGGUAGUAGGUUGUAUAGUU) with FAIMS NF directly sprayed into the MS with almost no spectra aside from contaminant single charged species. (Top) Spectra of 10nM Let7 microRNA with FAIMS attached at a CV of 55 with the distribution of charge states and each of them being nuclear envelopes of [biotin-Let7a]+2Na+H2O calculated to a reference. (Bottom) B) Comparison of FAIMS and NF with example oligonucleotide UCCCCCCCCUCGp with the sequence map coverage outlined in top right for both and a-d ions highlighted in green while w-z ions highlighted in red. C) Comparison of FAIMS and NF with example oligonucleotide UCCACUCAGp with the sequence map coverage outlined in top right for both and a-d ions highlighted in green while w-z ions highlighted in red.

**Supplemental Figure 2.**
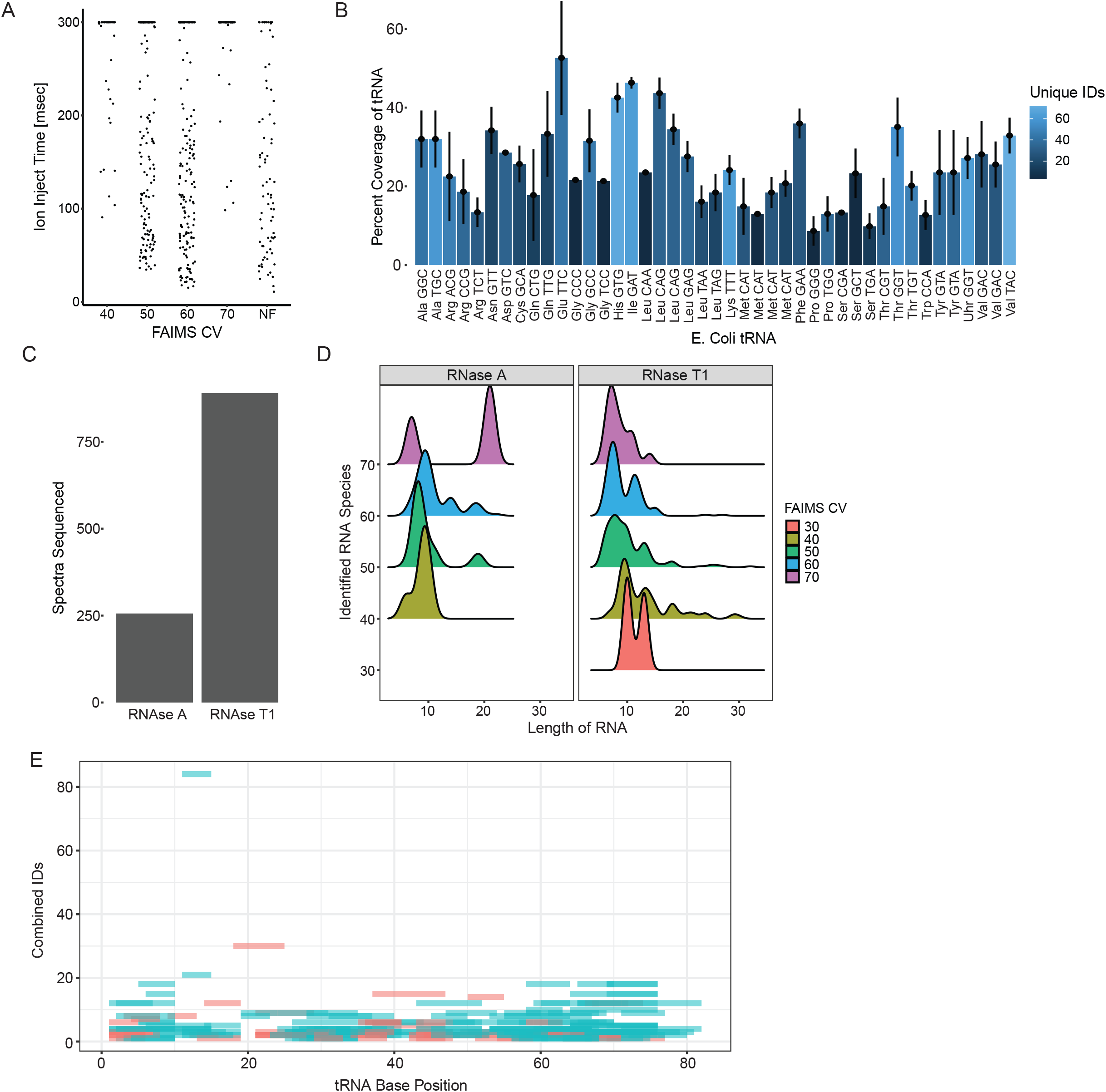
A)Ion Inject times of RNase T1 treated tRNA in different CV values in direct injection. B) Coverage of tRNA genes identified by FAIMS in triplicate technical replicates with unique spectra per gene identified by color bar. C) Direct Injection FAIMS comparing spectra sequenced from RNase A and RNase T1 digestion treatments. D) Distributions of sequenced RNA lengths between FAIMS CV and RNase treatment. E) Overlapping identifications of RNase treatment with RNase A (red) and RNase T1 (blue) showing individual sequence coverage across all tRNAs.

## Notes

### Competing Interest Statement

The authors have declared no competing interest.

